# *In silico* drug design of benzothiadiazine derivatives interacting with bilayer cell membranes

**DOI:** 10.1101/2022.02.14.480348

**Authors:** Zheyao Hu, Jordi Marti

## Abstract

The use of drugs derived from benzothiadiazine, a bicyclic heterocyclic benzene derivative, has become a widespread treatment for diseases such as hypertension, low blood sugar or the human immunodeficiency virus, among others. In this work we have investigated the interactions of benzothiadiazine and several selected derivatives designed *in silico*, with the basic components of cell membranes and solvents such as phospholipids, cholesterol and water. The analysis of the mutual microscopic interactions is of central importance to elucidate the local structure of benzothiadiazine as well as the mechanisms responsible for the distribution and access of benzothiadiazine to the interior of the cell. We have performed molecular dynamics simulations of benzothiadiazine and its derivatives embedded in a model zwitterionic bilayer membrane made by phospholipids dioleoylphosphatidylcholine, 1,2-dioleoyl-sn-glycero-3-phosphoserine and cholesterol inside aqueous potassium chloride solution in order to systematically examine microscopic interactions of benzothiadiazine derivatives with the cell membrane at liquid-crystalline phase conditions. From data obtained through radial distribution functions, time dependent hydrogen-bond lengths and potentials of mean force based on reversible work calculations, we have observed that benzothiadiazine derivatives have a strong affinity to stay at the cell membrane interface although their solvation characterisitics can vary significantly: they can be fully solvated by water in short periods of time or continuously attached to specific lipid sites during intervals of 10-70 ns. Furthermore, benzothiadiazines are able to bind lipids and cholesterol chains by means of single and double hydrogen-bonds of different characteristic lengths between 1.6 and 2.1 Å.

## 1. Introduction

Cell membranes are fundamental in the behavior of human cells, not only responsible for the interactions between the cell and its environment but also for processes such as cellular signaling[1], enzyme catalysis[2], endocytosis[3] and transport, among others. The main structure of the cell membrane is composed of bilayer phospholipids including sterols, proteins, glycolipids and a wide variety of other biological molecules. High compositional complexity and versatility of membranes are closely related to the environment and the physiological state of cells[4,5] so that many diseases such as cancer, cardiopathies, diabetes, atherosclerosis, infectious diseases or neurodegenerative pathologiesare accompanied by changes in the composition of cell membranes[6–9]. For such a reason, the knowledge of the behaviour of drugs interacting with different membrane components and their distribution in damaged tissues maybe key to improving drug efficiency and the therapy of the diseases and it has become a topic of greatest scientific interest.

It is well known that the composition of cell membranes in different tissues and organs of the human body shows large variations. In the treatment of diseases, an efficient drug design could enhance the interaction of active pharmaceutical ingredients with membrane components in specific tissues helping to reach the target site successfully. Thus, there is a great demand for a full understanding of the rules of drug-membrane interactions which may help us predict the distribution and curative effect of drugs in the body when it comes to the designing and testing of new drug molecules. Generally, medicinal chemists tended to overcome the difficulty of drugs in entering cells or crossing biological barriers, such as the blood-brain barrier[10–12] by modifying their structures to enhance the lipophilicity of drugs. However, little research has been done on the influence of drug structure on the rule of drug-membrane interaction, notably, the direct information on atomic interactions of drug-membrane systems at the all-atom level. In this work we will devote ourselves to establish a procedure for the *in silico* design of derivatives of the well-know family of benzothiadiazines.

Heterocyclic are ubiquitous in the structure of drug molecules[13,14] playing an important role in human life[15,16]. Such compounds are common parts of commercial drugs having multiple applications based on the control of lipophilicity, polarity, and molecular hydrogen bonding capacity. Among them benzothiadiazine and its derivatives have wide pharmacological applications, such as diuretic[17], antiviral[18], antiinflammatory[19], anticancer[20] and the regulation of the central nervous system[21], for instance. In addition to the above-mentioned biopharmacological activities, benzothiadiazine derivatives also has the bio-activity such as Factor Xa inhibition[22], anti-Mycobacterium[23,24], anti-benign prostatic hyperplasia[25]. 3,4-dihydro-1,2,4- benzothiadiazine-1,1-dioxide (DBD) being the main common structure of the benzothiadiazine family was investigated in a previous work[26] to elucidate the mechanisms responsible for the interactions of DBD with the basic components of cell membranes in all-atom level for the first time. The DBD has a strong affinity to the DOPC species of lipids, is also able to bind other membrane component by single and double hydrogenbonds. In this paper, we modified DBD and evaluate the effect of different substituents on the affinity of the DBD to cell membrane components.

## 2. Methods

Five models of lipid bilayer membranes in aqueous solution have been constructed using the CHARMM-GUI web-based tool[27,28]. The membrane components and the amount of particles of each class are as follows: all systems include one single DBD derivative, 112 neutral DOPC lipids, 28 DOPS associated with K^+^ (DOPS-K) lipids, 60 cholesterol molecules, 49 potassium ions, 21 chlorine ions and 10000 water molecules. The lipids have been distributed in two symmetric leaflets embedded inside an electrolyte potassium-chloride solution at 0.15 M concentration. We have considered five different setups, where only the benzothiadiazine derivative is different in each case. We considered a previously investigated[26] standard DBD species as the reference (DBD1) and four more DBD derivatives (DBD2, DBD3, DBD4 and DBD5), designed by ourselves using *in silico* techniques. The way how we designed the new DBD species is as follows: Medicinal chemists modify the chemical structure of the drug for the purpose of improving the therapeutic effect of the drug, reducing the toxic and side effects. The modification method depends on the structure of the drug. Generally, when doing the structure modification, the basic structure of the drug will remain unchanged and only some functional group will change. When the drug acts the binding methods of drug and receptor to form a reversible complex are generally by ionic bond, hydrogen bond or covalent bond. In our previous work[26] we observed that DBD can form hydrogen bonds (HB) and become absorbed by the cell membrane with DBD having strong affinity for DOPC. ‘H2’ and ‘H4’ sites of DBD are important for the formation of such HB with membrane components. The ‘R’ site (shown in Fig. 1) is very close to the ‘H2’ and ‘H4’ sites so that the size, electronegativity and other properties of the R substituent will affect the ability of ‘H2’/’H4’ to form hydrogen bonds with cell membrane components. So, with the tool of CHARMM-GUI platform “Ligand Reader & Modeler”, we introduced methyl, ethyl, fluorine and trifluoromethyl into this site in order to assess the effect of new drug structures on the behavior of DBD in cell membranes.

**Figure 1.**
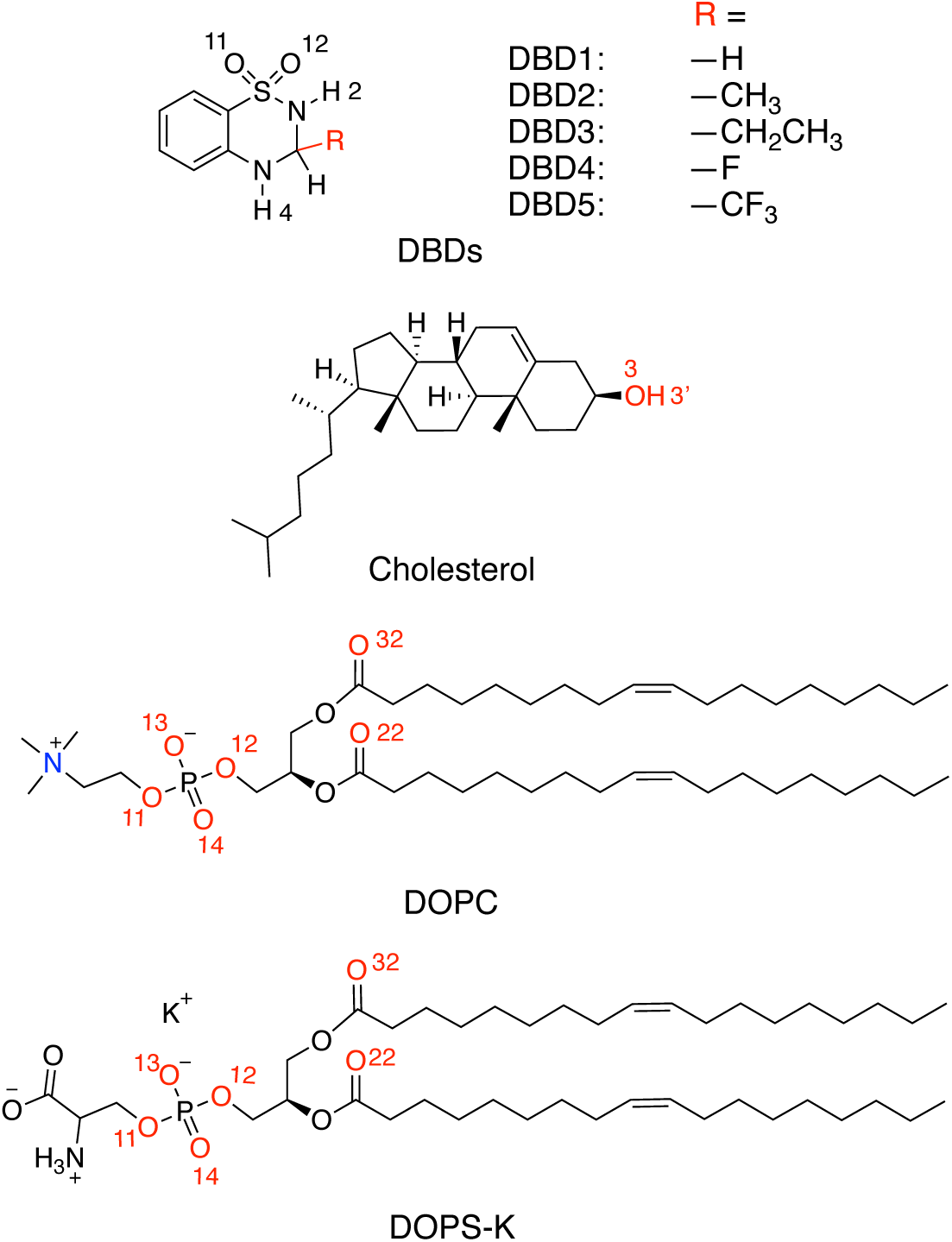
Chemical structures of benzothiadiazine derivatives, phospholipids and cholesterol. Site ‘R’ stands for the five DBD derivatives considered in the present work.

Sketches of all species are reported in Fig. 1. Each DBD species and phospholipid was described with atomic resolution (DBD1 and DBD4 have 20 sites, DBD2 and DBD5 have 23 sites, DBD3 has 26 sites, DOPC has 138 sites, DOPS has 131 sites and cholesterol has 74 sites). In all simulations water has been represented by rigid 3-site TIP3P[29] molecules. The CHARMM36 force field[30,31] was adopted for lipid–lipid and lipid–protein interactions. In particular, we selected the version CHARMM36m[32], which is able to reproduce the area per lipid for the most relevant phospholipid membranes, in excellent agreement with experimental data. The parameterization of the DBD species was performed by means of the “Ligand Reader & Modeler” tool in CHARMM- GUI platform (https://charmm-gui.org/?doc=input/ligandrm). All bonds involving hydrogens were set to fixed lengths, allowing fluctuations of bond distances and angles for the remaining atoms. Van der Waals interactions were cut off at 12 Å with a smooth switching function starting at 10 Å. Finally, long-ranged electrostatic forces were computed using the particle mesh Ewald method[33], with a grid space of about 1 Å, updating electrostatic interactions every time step of each simulation.

Molecular dynamics (MD) simulations have been revealed to be a very reliable tool for the simulation of the microscopic structure and dynamics of all sorts of condensed systems, such as aqueous solutions in bulk or under confinement[34–37] toward model cell membranes in electrolyte solution[38–40] and, more recently, small-molecule and protein systems attached to phospholipid membranes[41–43]. Five sets of MD runs were performed by means of the GROMACS2021 simulation package[44–48]. We run all the simulations at the fixed pressure of 1 atm and at the temperature of 310.15 K, typical of the human body and also well above the crossover temperatures for pure DOPC and DOPS needed to be at the crystal liquid phase (253 and 262 K, respectively)[49]. In all cases, the temperature was controlled by a Nose-Hoover thermostat[50] with a damping coefficient of 1 ps^−1^, whereas the pressure was controlled by a Parrinello-Rahman barostat[51] with a damping time of 5 ps. In the isobaric–isothermal ensemble, i.e., under the condition of a constant number of particles, pressure and temperature, equilibration periods for all simulations were around 1.875 ns. In all cases, we recorded statistically meaningful trajectories of 600 ns. The simulation boxes had same size in all cases, i.e. 78.1 × 78.1 × 95.7 Å^3^. We have considered periodic boundary conditions in the three directions of space. The simulation time step was fixed to 2 fs.

## 3. Results and Discussion

### 3.1 Characteristics of the bilayer systems

The phospholipid bilayer considered in this work was previously simulated and its main characteristics were reported[26,52]. We found reliable values of the area per lipid *A* and the thickness Δ*z* of the membranes to be in qualitative agreement with available experimental data. In order to corroborate these results in the present work where the system contains DBD derivatives, we computed *A* and Δ*z* as usual, considering the total surface along the *XY* plane (plane along the bilayer surface) divided by the number of lipids and cholesterol in one single leaflet[53] and the difference between the *z*-coordinates of the phosphorus atoms of the two leaflets, respectively. The results of the averaged values obtained from the 600 ns production runs are reported in Table 1, whereas the time evolution of both properties is displayed in Fig. 2.

**Table 1.** Physical characteristics of the systems simulated in this work. Area/lipid and thickness given in Å^2^ and Å units, respectively.

| DBD derivative | $A$ | $\Delta z$ |
| --- | --- | --- |
| DBD1 | 52.18 | 43.04 |
| DBD2 | 52.20 | 43.02 |
| DBD3 | 52.23 | 42.98 |
| DBD4 | 52.19 | 43.02 |
| DBD5 | 52.23 | 42.97 |

**Figure 2.**
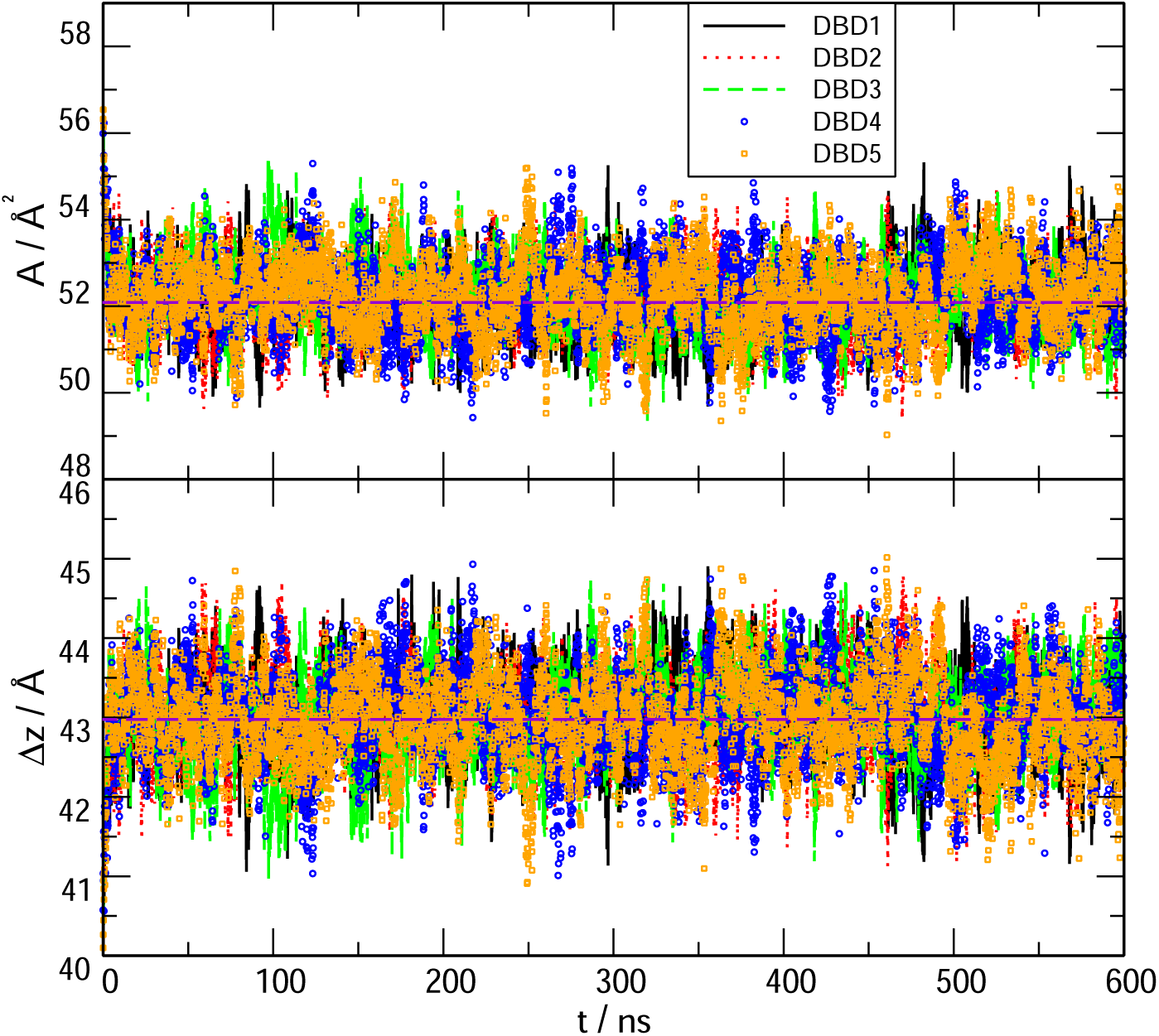
Area per lipid *A* and thickness Δ*z* of the membrane systems including DBD derivatives as a function of simulation time *t*. DBD1 (continuous line); DBD2 (dotted line); DBD3 (dashed line); DBD4 (circles); DBD5 (squares). Long-dashed (purple) lines indicate the average values reported in Table 1.

The results shown in Fig. 2 indicate that the simulated trajectories were well equilibrated in all cases. The comparison with previous results indicates that the effect of DBD derivatives on the area per lipid and thickness of the membrane is totally marginal. Firstly, the averaged result of *A* = 52.2 Å^2^ in all cases matches perfectly the previous reported value of 52.0 Å^2^[52] (where a large protein was embedded in the system) and also the experimental value of 54.4 Å^2^ reported by Nagle et al.[54]. Area/lipid shows fluctuations around 5% of the averaged values. Secondly, thickness of the membranes are also in good qualitative agreement with previous works: from Fig. 2 we observe fluctuations less than 5% of the averaged values, of around 43.0 Å, as expected. Such value is in qualitative agreement with the experimental measurement of Δ*z* = 40 Å for the DOPC- cholesterol (30%) bilayer, as reported by Nagle et al.[54] and it matches the previously found Δ*z* = 43.0 Å obtained in previous simulations for the DOPC/DOPS/cholesterol membrane[52].

### 3.2. Local structure of benzothiadiazine derivatives

#### 3.2.1. Radial distribution functions

We considered the so-called atomic pair radial distribution functions (RDF) *g*_*AB*_(*r*), defined, in a multicomponent system, for a species *B* close to a tagged species *A* as:

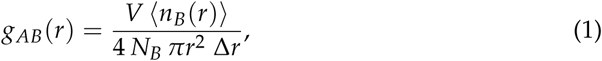

where *n*_*B*_(*r*) is the number of atoms of species *B* surrounding a given atom of species *A* inside a spherical shell of width Δ*r. V* is the total volume of the system and *N*_*B*_ is the total number of particles of species *B*. The physical meaning of the RDF stands for the probability of finding a particle *B* at a given distance *r* of a particle *A*. Our RDF are normalised so that tend to 1 at long distances, i.e. when the local density equals the averaged one.

We have evaluated the local structure of the DBD derivatives when solvated by lipids, cholesterol and water according to Eq.1. Only a few of all possible RDFs are reported, since we have selected the most relevant ones for the purpose of highlighting the main interactions between the tagged particles. The results are presented in Figures 3, 4 and 5, where we have selected the hydrogen sites ‘H2’, ‘H4’ and ‘O11-O12’ of DBD derivatives, since these are the most active sites, able to form hydrogen bonds (HB) with the surrounding partners (lipid, cholesterol species and eventually water). In all cases we can observe a clear first coordination shell associated to the binding of DBD derivatives to the membranes, with corresponding maxima indicating the typical HB distances, together with much lower second shells centred around 4-5 Å. As a general fact, the HB detected cover a noticeably wide range of distances, between 1.6 and 2.1 Å.

**Figure 3.**
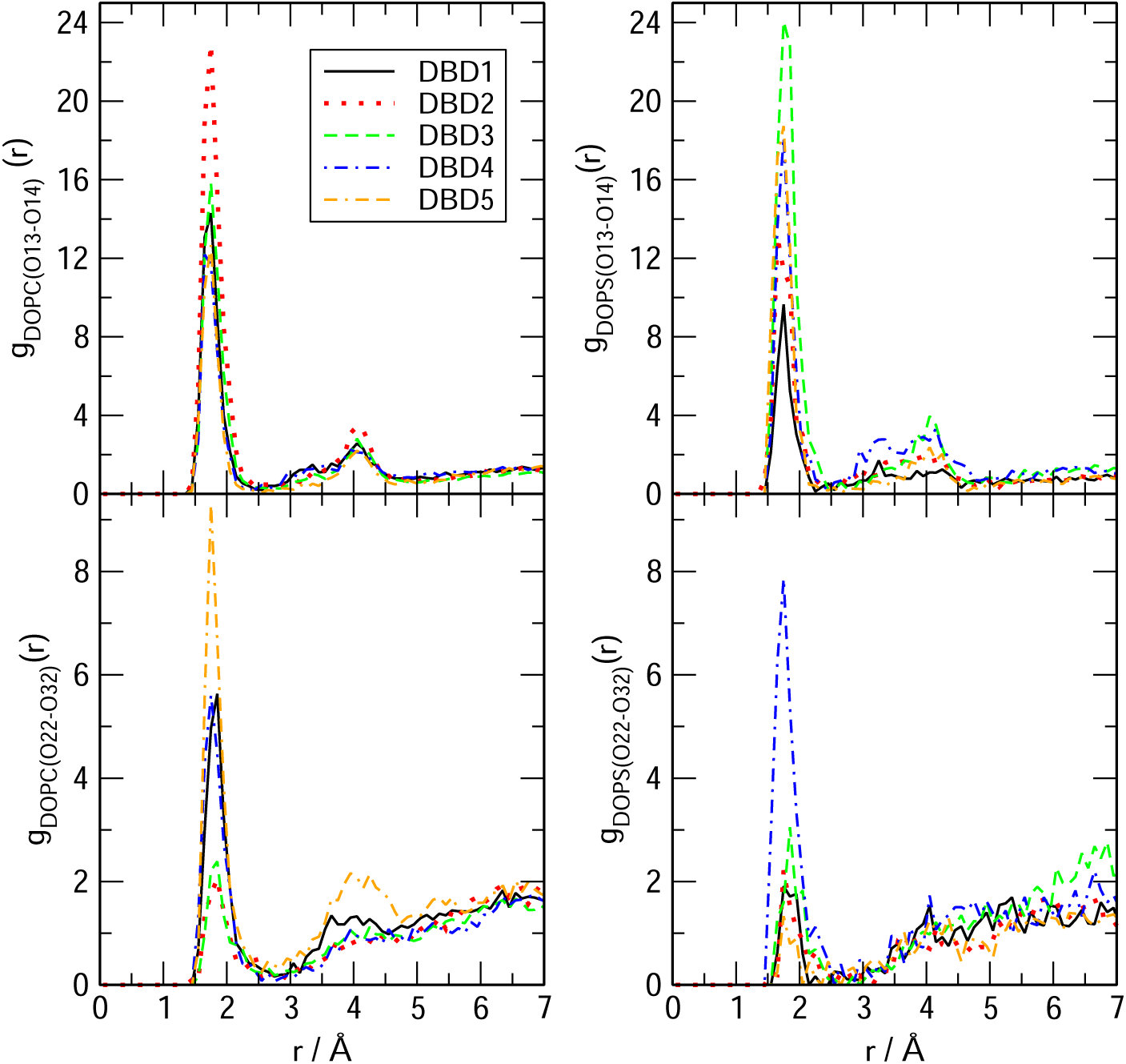
Radial distribution functions between site ‘H2’ of DBD derivatives and selected oxygen sites in DOPC and DOPS phsopholipids. Sites ‘O13-14’ stand for head-groups of the cell membrane phospholipids and sites ‘O22-32’ stand for tail-groups located deeper in the membrane interface.

**Figure 4.**
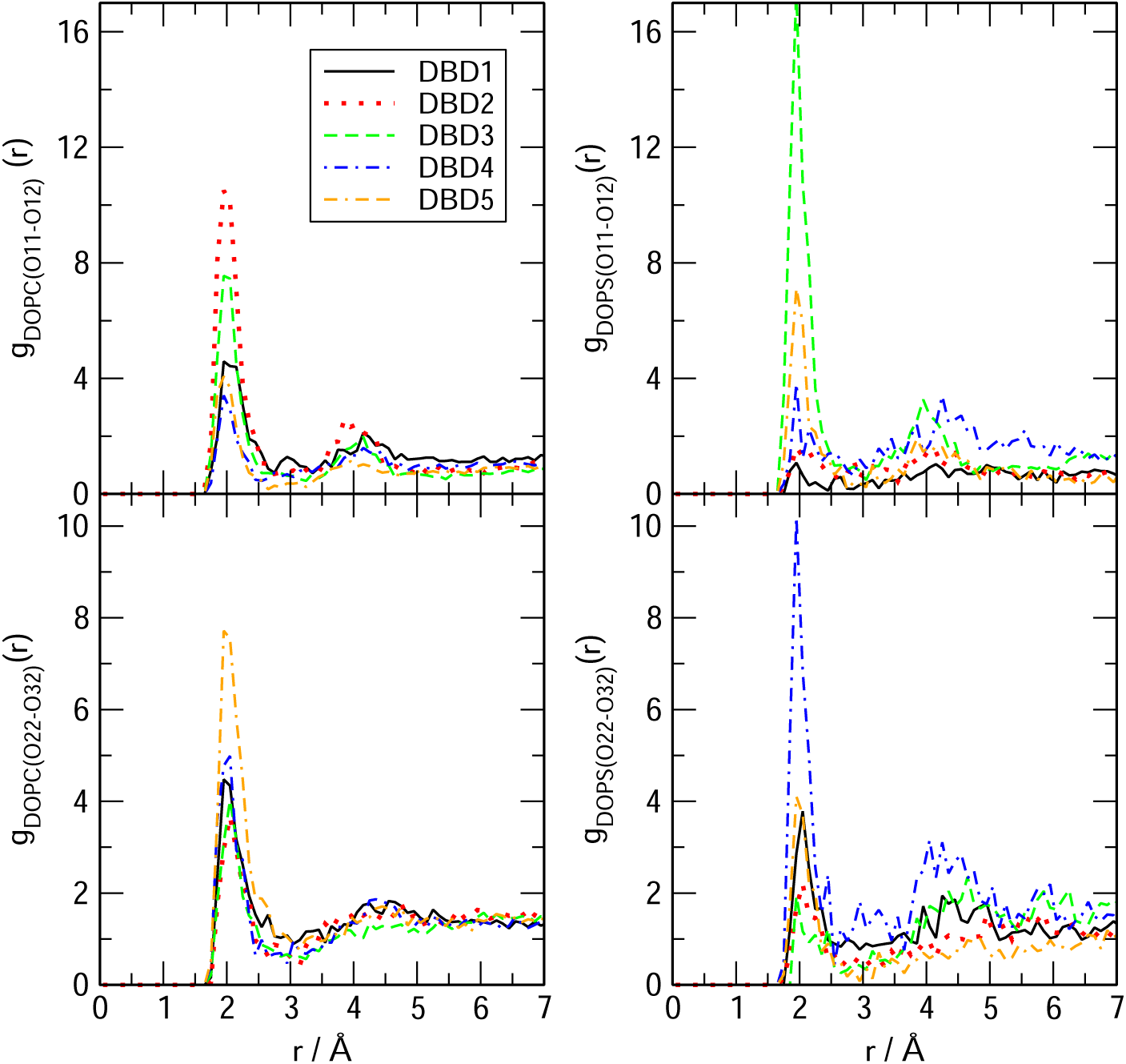
Radial distribution functions between site ‘H4’ of DBD derivatives and selected oxygen sites in DOPC and DOPS phsopholipids.

**Figure 5.**
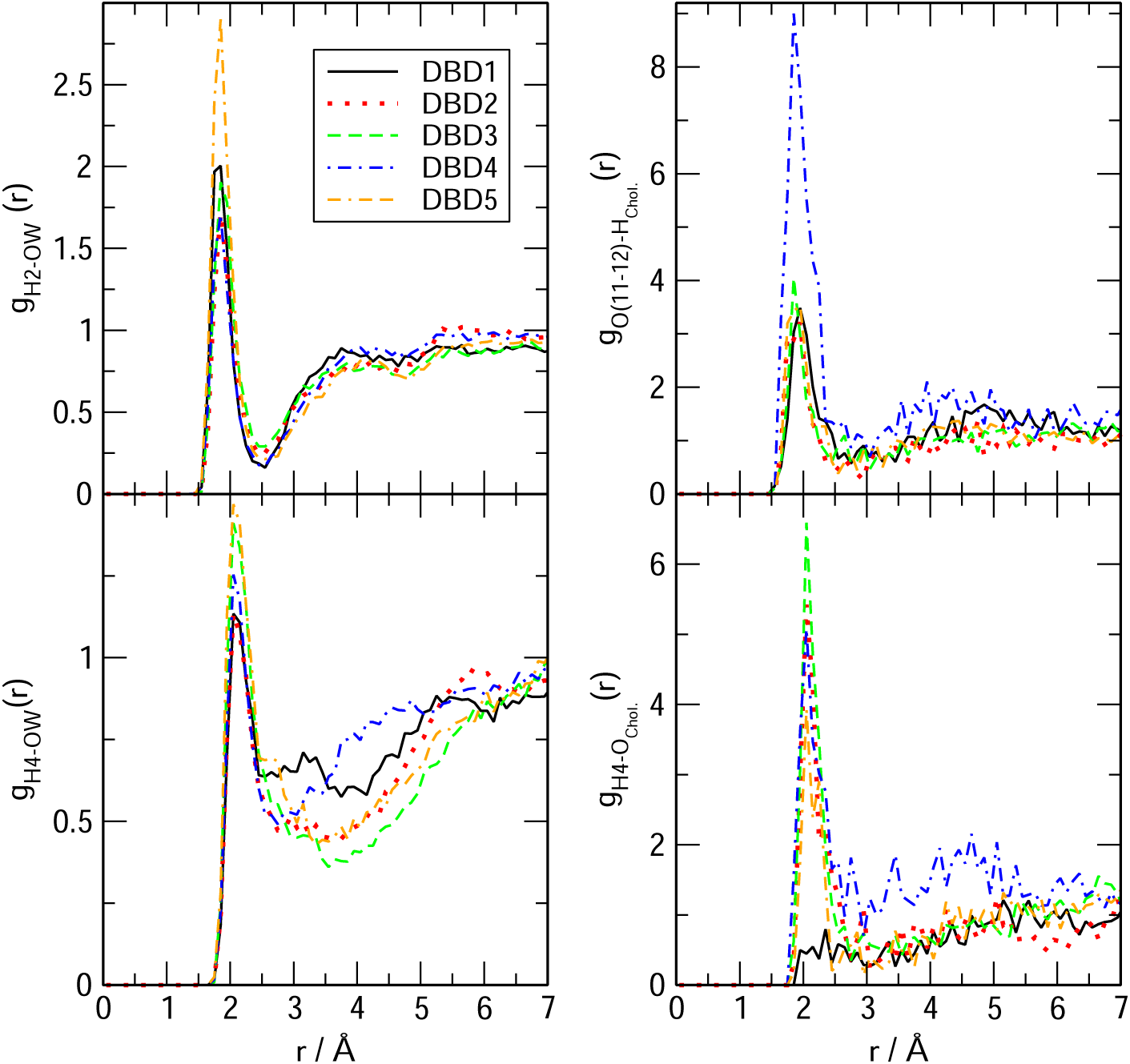
Radial distribution functions between sites ‘H2’ and ‘H4’ of DBD derivatives and selected sites of water (left column) and cholesterol (right column).

The structure of DBD derivatives described by their ‘H2’ site (see Fig. 1) indicates the existence of HB formed by ‘H2’ and several sorts of lipid oxygen sites and it is represented in Fig. 3. We can notice that the typical HB length is of 1.7 Å in all cases, both for the binding with oxygens of the phosphoryl group ‘O13-14’ (located at the head-groups of DOPC and DOPS, with both oxygen sites sharing a negative charge) and for the binding with sites ‘O22-32’ (located in the tail-groups of the lipids) as well. This is the typical distance of the binding of small-molecules to cell membranes, such as tryptophan to dipalmytoilphosphatidylcholine (see for instance the review [55]). It should be pointed out that using fluorescence spectroscopy, Liu et al.[56] obtained values for the HB lengths of tryptophan-water between 1.6 and 2.1 Å, i.e of the same range than those reported here.

This indicates that: (1) all sorts of DBD derivatives can bind the membranes in both head- and tail-groups and (2) depending on the oxygen sites, some derivatives are able to create HB stronger than others. However, the strength of the HB binding is not uniform and it clearly depends of the class of derivative and lipid chain involved. Despite we will qualitatively analyze the strength of the HB in Section 3.2.2, we can give some general clues. For instance, DBD2 is able to bind DOPC more strongly than DBD1 (reference), with the remaining derivatives making bonds of similar strength. Nevertheless, when DOPS is concerned, all derivatives form stronger HB than DBD1, with DBD3 the strongest. Similar trends are observed when the internal tail-group sites ‘O22-23’ are analized: DBD5 makes the strongest HB with DOPC and DBD4 makes the strongest bond with DOPS. In this latter case, the enhancement of the HB is milder than it occured in the former case (head-group bindings). As a general fact, we can indicate that the one-site modifications proposed with the design of the new DBD-derivatives reported in this work has produced significant changes and enhancement of the HB connections to the model cell membrane. This can be valuable information to assess the affinity of new designed drugs to target specific oncogenes such as KRas-4B, work that it is been currently developed in our group.

Concerning hydrogens ‘H4’ of DBD derivatives and their binding characteristics when associated to DOPC and DOPS (Fig. 4), we observed that they can be also connected either to ‘O11’ or ‘O12’ of the phospholipids (head-groups), either to ‘O22-O32’ (tail-groups). Interestingly, in the case of DBD’s ‘H4’, HB lengths are within the range of 2-2.1 Å, significantly longer than those formed by H2 (range around 1.6 to 1.8 Å). This was already observed for the reference DBD1 in a previous work[26]. In this case, the strongest HB is observed when ‘H4’ of DBD3 is connected to DOPS’s ‘O11-12’ oxygens. In this particular case, the new DBD derivatives have shown to be able to bind the internal regions of the membrane, whereas the original benzothiadiazine species (DBD1) had a very low probability to penetrate these regions. Again for the ‘H4’ binding site, we have found a general enhancement of the binding of DBD derivatives with the main phospholipids forming our cell membrane system.

In the third RDF set (Fig. 5) we report interactions between sites ‘H2’, ‘H4’ and ‘O11-12’ of DBD with water (plots at the left column) and cholesterol (plots at the right column). In the case of water, HB can be established between ‘H2’ and the oxygen site of water (top plot) or, alternatively, between ‘H4’ and the oxygen of water (bottom plot). In both cases, the strength of the interaction is low, what suggests that DBD derivatives are strongly bound to the cell membrane and can be solvated by a few water molecules located at the interface. We have not observed long term episodes of DBD derivatives fully solvated by water.

We have located some extent of hydrogen-bonding between DBD and cholesterol. However, no significant binding of ‘H2’ with cholesterol has been observed, whereas interactions of both ‘H4’ and ‘O11-12’ sites of DBD derivatives have been detected. In particular, the strongest contributions are seen for with hydroxyl’s oxygens of cholesterol with ‘H4’ of the DBD species, which were undetected for the reference original DBD1 as well as for oxygens of the DBD derivatives with hydroxyl’s hydrogen of cholesterol. In the latter case, we found a particularly strong contribution of DBD4, i.e. the derivative containing a fluoride residue instead the original hydrogen atom. The HB lengths are in the range of 2.1 Å in all cases.

#### 3.2.2. Potentials of mean force between benzothiadiazine derivatives and lipids

Among the wide variety of one-dimensional free-energy methods proposed to compute the potential of mean force (PMF) between two tagged particles[57] a simple but meaningful choice is to consider the radial distance *r* as an order parameter, able to play the role of the reaction coordinate of the process, within the framework of unbiased simulations as those reported in the present work and to proceed with a direct estimation of the reversible work as it will be described below. This has become one of standard choices to compute free-energy barriers in MD simulations, together with constrained MD simulations[58] or the popular *umbrella sampling* procedure[59]. In case that more accurate values of the free-energy barriers are needed, the optimal choices are: (1) to use constraint-bias simulation combined with force averaging for Cartesian or internal degrees of freedom[57]; (2) the use of multi-dimensional reaction coordinates[60] such as transition path sampling[61–63] or (3) considering collective variables, such as metadynamics[52,64] although such methods require a huge amount of computational time. Since the determination of reaction coordinates for the binding of DBD at zwitterionic membranes is out of the scope of this work, we will limit ourselves to use radial distances between two species as our order parameters to perform reversible work calculations.

In this framework, a good approximation of the PMF can be obtained by means of the reversible work *W*_*AB*_(*r*) required to move two tagged particles (A,B) from infinite separation to a relative separation *r* (see for instance Ref.[65], chapter 7):

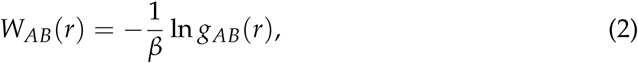

where *β* = 1/(*k*_*B*_*T*) is the Boltzmann factor, *k*_*B*_ the Boltzmann constant and T the temperature. In the calculations reported here, the radial distance *r* is the distance used in the corresponding RDFs (Section 3.2)i.e. it is not related to the atom position relative to the center of the membrane. All free-energy barrier are simply defined (in *k*_*B*_*T* units) by a neat first minimum and a first maximum of each *W*(*r*), with barrier size Δ*W* obtained as the difference between the former. As a sort of example, we present the free-energy barriers with largest values for each DBD species in Fig. 6.

**Figure 6.**
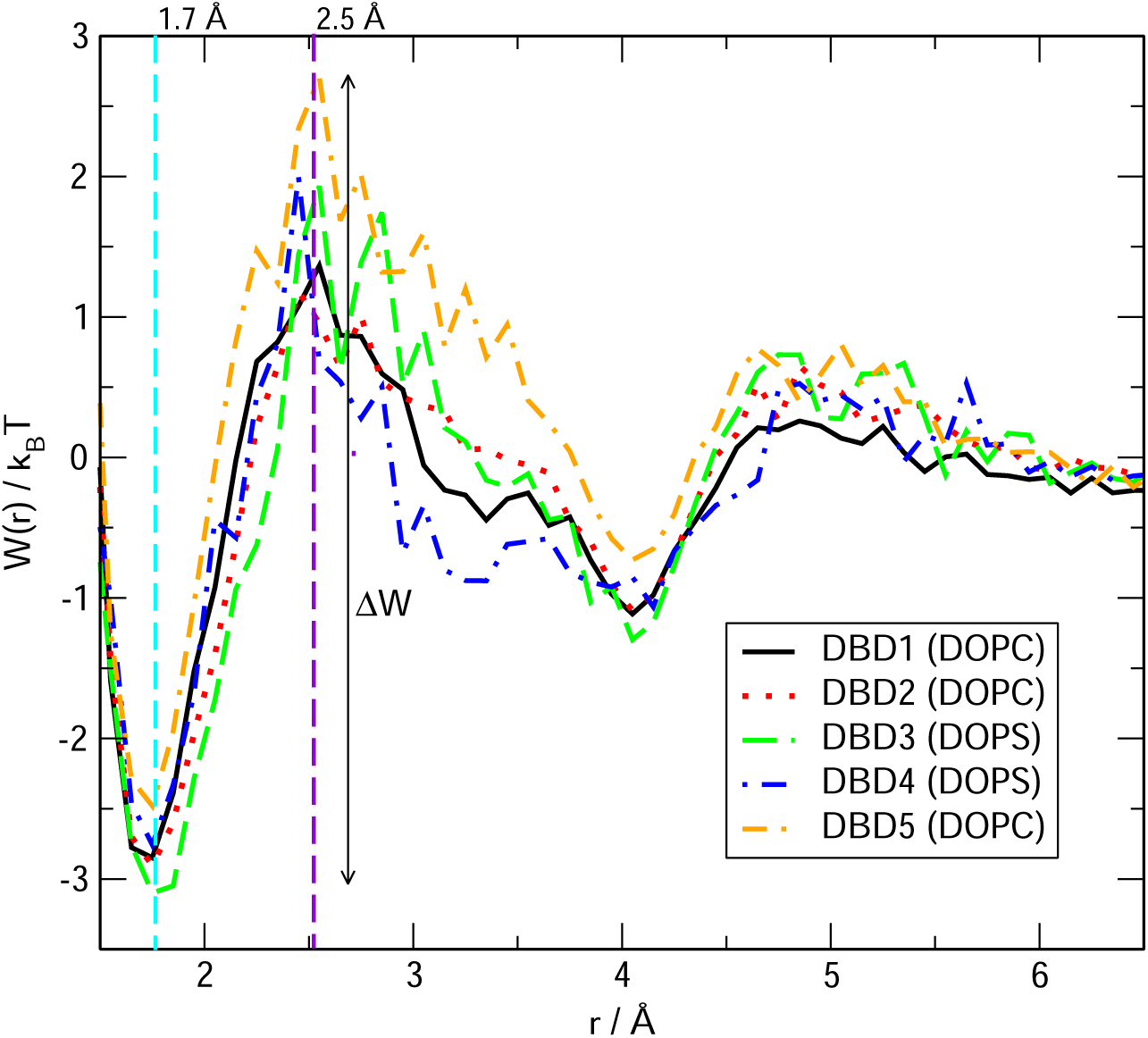
Potentials of mean force for the binding of ‘H2’ sites of DBD derivatives to the sites ‘O13-O14’ of DOPC and DOPS.

The full set of free-energy barriers for a wide selection of bound pairs has been reported in Table 2. There we can observe overall barriers between 1.2 and 5.2 *k*_*B*_*T*, what correspond to 0.7-3.1 kcal/mol, for the simulated temperature of 310.15 K. We observe stable binding distances (given by the position of the first minima of the PMF) matching the typical hydrogen-bond distances, as expected. As a reference, it is known that the typical energy of water-water hydrogen-bonds estimated from *ab-initio* calculations is of 4.9 kcal/mol for a water dimer in vacuum[66], whereas in our model system (including TIP3P water) the barrier associated to the HB signature, given by the first maximum of water’s oxygen-hydrogen RDF, is of 1.1 kcal/mol. This low value can be directly associated with two facts: (1) first, we have estimated this energy in the bulk, condensed phase of the aqueous ionic solvent, whereas the reference value of Feyereisen et al.[66] corresponds to an isolated water dimer, i.e. can be related to gas phase; (2) secondly, the TIP3P water model included in the CHARMM36 force field is well known to have significant failure to describe liquid water[67].

**Table 2.** Free-energy barriers ΔW (in *k*_*B*_*T*) from reversible work calculations for the binding of DBD to cholesterol, lipids and water. In order to quantify the height of all barriers, 1 *k*_*B*_*T* = 0.596 kcal/mol. Labels as indicated in Figure 1.

| DBD site | Lipid site | $\Delta W$ | | | | |
| --- | --- | --- | --- | --- | --- | --- |
|  |  | DBD1 | DBD2 | DBD3 | DBD4 | DBD5 |
| H2 | O13-O14 DOPC | 4.2 | 4.0 | 4.0 | 4.2 | 5.2 |
| H2 | O22-O32 DOPC | 3.4 | 3.2 | 2.5 | 4.4 | 3.7 |
| H2 | O13-O14 DOPS | 4.2 | 3.8 | 5.0 | 4.7 | 4.7 |
| H2 | O22-O32 DOPS | 2.8 | 3.5 | 3.3 | 3.0 | 2.2 |
| H4 | O11 DOPC | 2.0 | 2.5 | 2.4 | 1.9 | 3.1 |
| H4 | O22-O32 DOPC | 1.6 | 2.2 | 1.9 | 2.3 | 2.3 |
| H4 | O11 DOPS | 2.3 | 1.2 | 3.5 | 1.8 | 3.8 |
| H4 | O22-O32 DOPS | 1.8 | 1.6 | 2.0 | 4.0 | 3.9 |
| H4 | O Cholesterol | 1.2 | 3.2 | 2.9 | 2.4 | 3.4 |
| O11 -O12 | H Cholesterol | 1.8 | 2.4 | 2.2 | 2.6 | 2.3 |

In our earlier work[26] we reported by the first time DBD-membrane related freeenergy barriers. For the sake of comparison with other similar systems, we can report that the PMF of tryptophan in a di-oleoyl-phosphatidyl-choline bilayer membrane shows a barrier of the order of 4 kcal/mol[68], whereas the barrier for the movement of tryptophan attached to a poly-leucine *α*-helix inside a DPPC membrane was reported to be of 3 kcal/mol[69]. Finally, neurotransmitters such as glycine, acetylcholine or glutamate were reported to show small barriers of about 0.5-1.2 kcal/mol when located close to the lipid glycerol backbone[70]. These values could further indicate that our estimations match well the order of magnitude of the free-energy barriers for other small-molecules of similar size.

We designed two sets of DBD derivatives according to their characteristics: in DBD2 and DBD3 we replaced a hydrogen by electron-donating groups (methyl and ethyl, respectively) whereas in DBD4 and DBD5 we replaced a hydrogen by electron-accepting groups (fluorine and trifluoromethyl, respectively). Regardless of the type of replacement considered, our general result is that most of the barriers are in the range of 1-5 kcal/mol, regardless of the specific derivative considered. As more specific features, we can observe that the barriers corresponding to the HB formed by the residue ‘H2’ of the DBD derivatives are overall larger than those related to the hydrogen-bonds formed by ‘H4’, what suggests that ‘H2’ is the most stable binding site between DBD species and the model cell membranes considered in this work. Among the five DBD species analysed we can observed that, regarding the ‘H2’ site of DBD, interactions of its derivatives with DOPC are about 10% stronger that those with DOPS but when ‘H4’ is concerned, the strength of its HB with DOPC is weaker than those with DOPS only when the tail-groups ‘O22-32’ are considered. Nevertheless, the barriers of ‘H4’ to head-groups are of similar size for both DOPC and DOPS. Finally, the binding of DBD with cholesterol is revealed to be sensibly weaker than that to DOPC and DOPS.

With the aim of a better understanding of the geometrical shape of the HB established between DBD and lipid species, we report in Fig. 7 a series of three snapshots describing the simultaneous binding of DBD4 with a few counterparts: so, we can observe that DBD4’s ‘H2’ and ‘H4’ are able to bridge oxygens ‘O13’ and ‘O14’ of DOPC and ‘O22-O32’ of DOPS (A), also ‘O22-O32’ of DOPC and ‘O’ of the hydroxyl group of cholesterol and finally ‘O13’ and ‘O14’ of DOPS and ‘O’ of the hydroxyl group of cholesterol. This remarkable bridging properties of DBD4 are qualitatively similar to those of DBD1. Both species, and to some extent all of DBD derivatives, can also form closed-ring structures (see Ref.[26], Figure 6). The bridging bonds highlighted here are quite similar to the HB structures observed in tryptophan[71] and melatonin absorbed at cell membrane surfaces[72].

**Figure 7.**
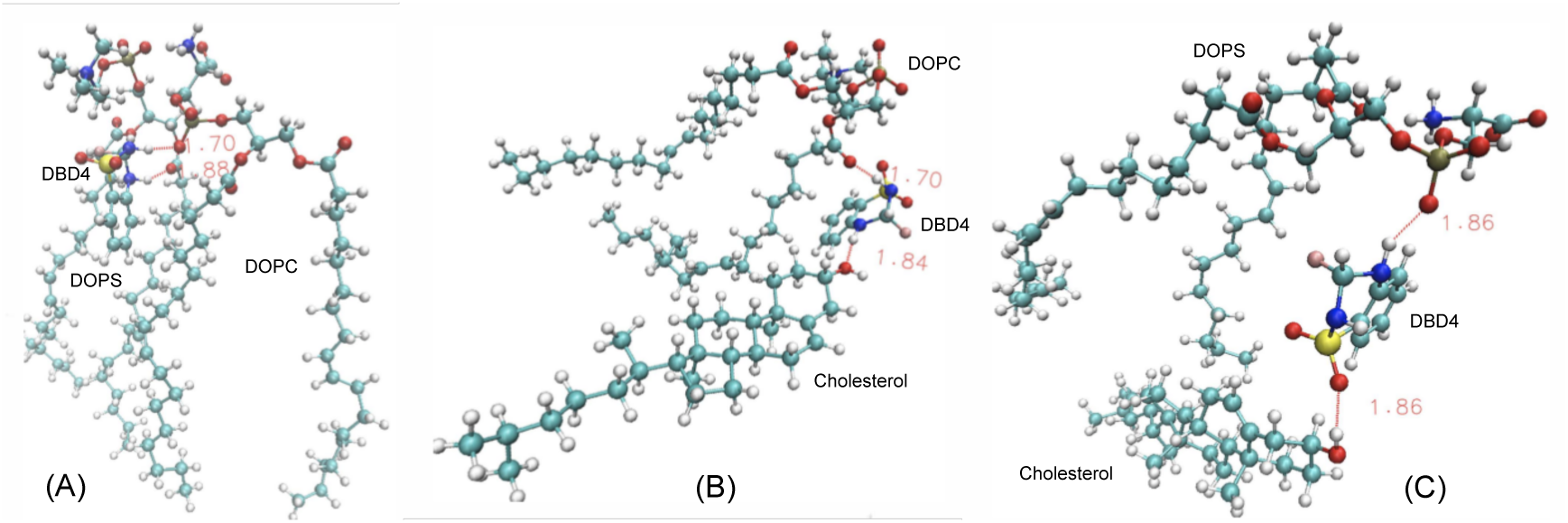
Snaphsots of relevant configurations between benzothiadiazine derivative DBD4 and their partner hydrogen-bonding sites. We indicate the site belonging to DBD4 in each case: (A) Site ‘H2’ bridging one DOPC lipid and site ‘H4’ bridging one DOPS lipid; (B) site ‘H2’ bridging one DOPC lipid and site ‘H4’ bridging one cholesterol molecule; (C) site ‘H4’ bridging one DOPS and ‘O’ site bridging one cholesterol molecule. Typical hydrogen-bond distances are indicated in red.

#### 3.2.3. Dynamical atomic site-site distances

Once the local structures of the DBD derivatives have been fully evaluated, we will make an estimation of the HB dynamics by computing the average lifetime of some of the HB reported by RDF. Other typical MD properties involving time-correlation functions such as power spectra[73,74], relaxation times or self-diffusion coefficients[75,76] that were considered in previous studies, are out of the scope of this paper and have not been considered here. We display the time evolution of selected atom-atom distances *d*(*t*) in Figure 8 only for the pairings of ‘H2’ of DBD3 and sites ‘O13’ and ‘O14’ of DOPC and DOPS (top panel) and for ‘H4’ of DBD3 and sites ‘O11’ and ‘O12’ of DOPC and DOPS (bottom panel), as a sort of example. The full set of averaged values are reported in Table 3. We have selected in Fig. 8 representative intervals (of more than 100 ns) from the full MD trajectory of 600 ns where the pattern of formation and breaking of HB is clearly seen, including a large extent of fluctuations. This means that such patterns have been systematically observed throughout the whole trajectory.

**Table 3.** Averaged distances (in Å) between selected sites of DBD and the membrane. Continuous time intervals (*τ*, in ns) have been obtained from averaged computations along the 600 ns trajectory. Labels as indicated in Figure 1.

| DBD site | Lipid site | Distance | $\tau_{DBD1}$ | $\tau_{DBD2}$ | $\tau_{DBD3}$ | $\tau_{DBD4}$ | $\tau_{DBD5}$ |
| --- | --- | --- | --- | --- | --- | --- | --- |
| H2 | O13-O14 DOPC | 1.7 | 67.4 | 73.7 | 63.2 | 28.4 | 38.5 |
| H2 | O22-O32 DOPC | 1.8 | 20.4 | 11.8 | 12.3 | 19.4 | 24.6 |
| H2 | O13-O14 DOPS | 1.7 | 6.9 | 8.8 | 27.0 | 9.6 | 9.7 |
| H2 | O22-O32 DOPS | 1.8 | 1.4 | 1.9 | 2.3 | 3.1 | 0.9 |
| H4 | O11-O12 DOPC | 1.9 | 42.5 | 64.4 | 61.5 | 15.6 | 35.9 |
| H4 | O22-O32 DOPC | 2.0 | 16.5 | 14.0 | 13.0 | 16.6 | 28.4 |
| H4 | O11-O12 DOPS | 1.9 | 1.3 | 1.9 | 24.8 | 1.9 | 6.4 |
| H4 | O22-O32 DOPS | 2.0 | 1.8 | 1.4 | 0.9 | 2.6 | 1.9 |
| H4 | O Cholesterol | 2.0 | 0.5 | 3.4 | 4.4 | 3.7 | 3.6 |
| O11-O12 | H Cholesterol | 1.9 | 4.6 | 4.5 | 6.6 | 10.3 | 6.0 |

**Figure 8.**
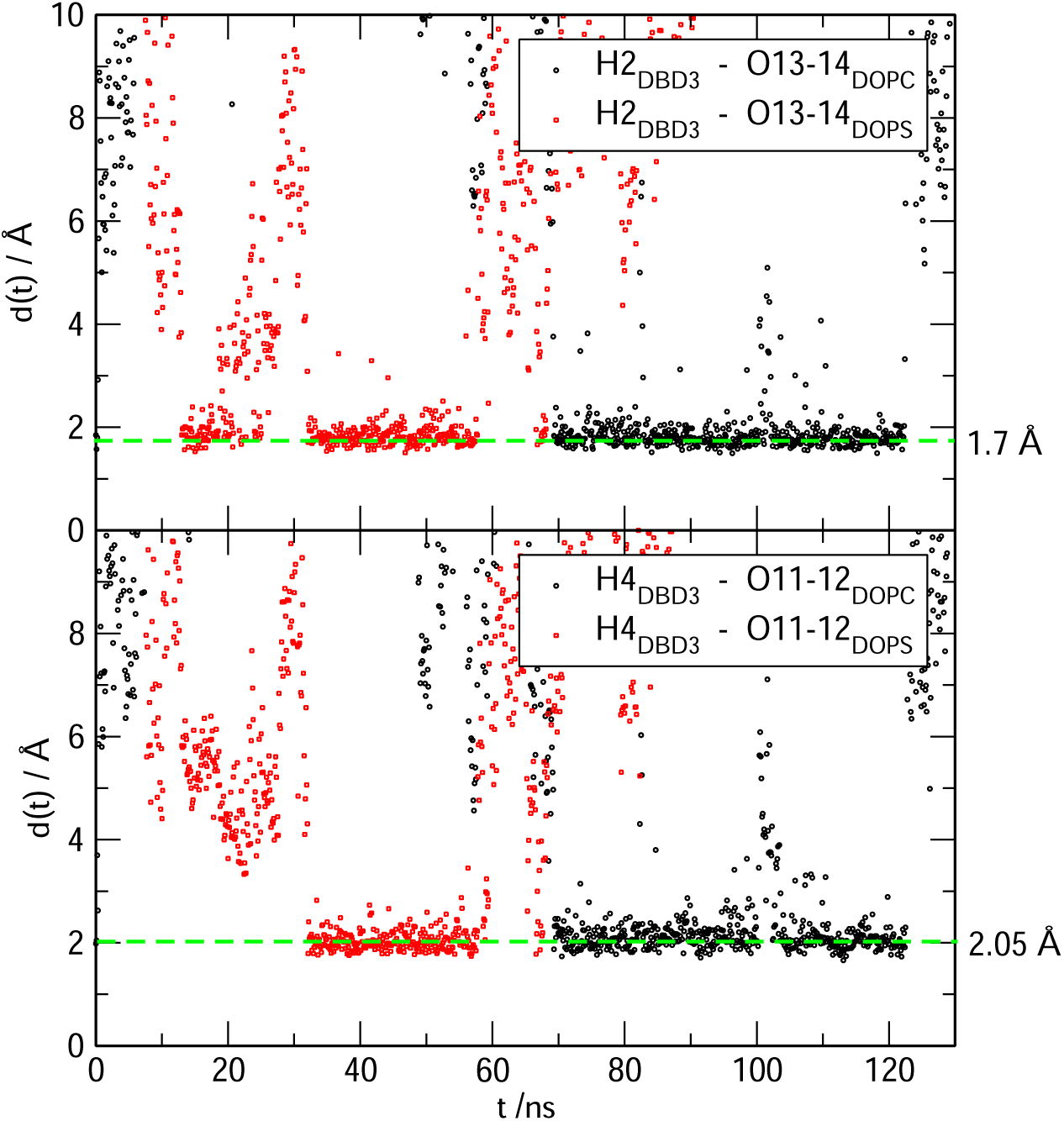
Time evolution of distances between selected sites of DBD3 (‘H2’, ‘H4’) and their partner oxygen sites of DOPC and DOPS.

We can observe that typical HB distances of 1.7 and 2.05 Å are reached. Sites ‘O13’ and ‘O14’ (and ‘O11’ and ‘O12’) of DOPC and DOPS have been averaged given their equivalence. Typical HB lifetimes can vary enormously, between short lived HB of less than 1 ns (DBD1 with cholesterol) up to long-life HB of more than 70 ns (DBD2 with the head-group of DOPC, i.e. sites ‘O13-O14’). As general trends, we can highlight that (1) sites ‘H2’ of the benzothiadiazine derivatives are able to form much longer lived HB than sites ‘H4’, especially for the DOPS species and (2) DBD-cholesterol hydrogen bonds have rather short lifetimes in the range of 1-10 ns. A closer look indicates that the longest living HB established between DBD and cholesterol are those composed by cholesterol’s hydrogen as donor and oxygens of DBD as acceptors, about twice longer that HB formed by hydroxil’s oxygen of cholesterol and hydrogen ‘H4’ of DBD derivatives. For the sake of comparison, we should remark that the typical lifetime of hydrogen-bonds in pure water has been estimated to be of the order of 1 ps[77].

## 4. Conclusions

We report results from molecular dynamics simulations of benzothiadiazine derivatives embedded in a phospholipid bilayer membrane formed by 200 lipid with 56% of DOPC, 14% of DOPS and 30% of cholesterol in aqueous potassium chloride solution using the CHARMM36m force field. Starting from a standard 3,4-dihydro-1,2,4- benzothiadiazine-1,1-dioxide molecule we have designed *in silico* four derivatives based on the replacement of a single hydrogen atom by two different classes of chemical groups, one of them electron-donating groups (methyl and ethyl) and another one by electron-accepting groups (fluorine and trifluoromethyl). As a gross feature, the same class of chemical groups produce similar effects on the HB between DBD and cell membranes, whereas different types tend to produce overall opposite effects. With this kind of study our aim is to ellucidate the effects of different chemical groups on DBD-cell interactions. Our analysis is based on the computation of the local structures of the DBD derivatives when associated to lipids, water and cholesterol molecules. After the systematic analysis of meaningful data, we have found that the location of DBD at the interface of the membrane is permanent. We have computed RDF defined for the most reactive particles, especially hydrogens ‘H2’ and ‘H4’ and oxygens ‘O11-O12’ of DBD (see Figure 1) correlated with sites of lipids and cholesterol able to form HB with DBD. All RDF have shown a strong first coordination shell and a weak second coordination shell for all DBD-lipid structures. The first shell is the signature of HB of lengths between 1.7 and 2.1 Å, in overall good agreement with experimental measurements[56] for comparable small-molecules at interfacial membranes.

The analysis of PMF of DBD-lipid interactions has revealed free-energy barriers of the order of 1-3 kcal/mol (Table 2), with the largest barriers corresponding to hydrogenbonds between DBD’s ‘H2’ site and oxygens sites of DOPC and DOPS. However, it has been observed that DBD derivatives are able to bind to cholesterol as well as the two classes of phospholipids, providing bridging connections that are able to locally estabilize and compactify the cell membrane, although the area per lipid and thickness of the whole membrane are not affected by the presence of the DBD species in any case. The influence of cholesterol has been especially noted in the weakening of DBD-lipid HB connections what should be taken in consideration for the interaction of drugs with cell membranes from a pharmaceutical point of view. After a thorough analysis monitoring relative distances between tagged sites of DBD and lipids we have estimated the lifetime of HB by averaging data from the 600 ns MD trajectories to range in between 1 and 70 ns.

## Author Contributions

Conceptualization, Z.H. and J.M.; methodology, Z.H. and J.M.; validation, Z.H. and J.M.; formal analysis, Z.H. and J.M.; investigation, Z.H. and J.M.; resources, Z.H. and J.M.; data curation, Z.H.; writing—original draft preparation, Z.H. and J.M.; writing—review and editing, Z.H. and J.M.; visualization, Z.H. and J.M.; supervision, J.M.; project administration, J.M.; funding acquisition, J.M. All authors have read and agreed to the published version of the manuscript.

## Acknowledgments

We thank Dr. Huixia Lu for fruitful discussions and technical support.

## References

1. Escribá P.V., Sastre M., García-Sevilla J. A. Disruption of cellular signaling pathways by daunomycin through destabilization of nonlamellar membrane structures. Proc. Natl. Acad. Sci. 1995, 92, 7595.

2. Tong S., Lin Y., Lu S., Wang M., Bogdanov M., Zheng L. Structural Insight into Substrate Selection and Catalysis of Lipid Phosphate Phosphatase PgpB in the Cell Membrane. J. Biol Chem. 2016, 291, 18342.

3. Doherty G. J., McMahon H. T. Mechanisms of endocytosis. Annu. Rev. Biochem. 2009, 78, 857.

4. Escribá P.V., González-Ros J. M., Goñi F.M., Kinnunen P. K., Vigh L., Sánchez-Magraner L., Fernández A. M., Busquets X., Horvath I., Barceló-Coblijn G. Membranes: a meeting point for lipids, proteins and therapies. J. Cell Mol. Med. 2008, 12, 829.

5. Noutsi P., Gratton E., Chaieb S. Assessment of Membrane Fluidity Fluctuations during Cellular Development Reveals Time and Cell Type Specificity. PLoS One. 2016, 11.

6. Hakomori S. Aberrant glycosylation in cancer cell membranes as focused on glycolipids: overview and perspectives. Cancer Res. 1985, 45, 2405.

7. Simons K., Ehehalt R. Cholesterol, lipid rafts, and disease. J. Clin Invest. 2002, 110, 597.

8. Vigh L., Escribá P.V., Sonnleitner A., Sonnleitner M., Piotto S., Maresca B., Horvath I., Harwood J. L. The significance of lipid composition for membrane activity: new concepts and ways of assessing function. Prog. Lipid Res. 2005, 44, 303.

9. Kim Y., Shanta S. R., Zhou L. H., Kim K. P. Mass spectrometry based cellular phosphoinositides profiling and phospholipid analysis: a brief review. Exp. Mol. Med. 2010, 42, 1.

10. Jolliet-Riant P., Tillement J. P. Drug transfer across the blood-brain barrier and improvement of brain delivery. Fundam. Clin. Pharmacol. 1999, 13, 16.

11. Bodor N., Buchwald P. Barriers to remember: brain-targeting chemical delivery systems and Alzheimer’s disease. Drug Discov. Today. 2002, 7, 766.

12. Waterhouse R. N. Determination of lipophilicity and its use as a predictor of blood-brain barrier penetration of molecular imaging agents. Mol. Imaging Biol. 2003, 5, 376.

13. Rees C.W. Polysulfur-Nitrogen Heterocyclic Chemistry. Journal of Heterocyclic Chemistry. 1992, 29, 639.

14. Vitaku E., Smith D. T., Njardarson J. T. Analysis of the structural diversity, substitution patterns, and frequency of nitrogen heterocycles among U.S. FDA approved pharmaceuticals. J. Med. Chem. 2014, 57, 10257.

15. Horton D. A., Bourne G. T., Smythe M. L. The combinatorial synthesis of bicyclic privileged structures or privileged substructures. Chem. Rev. 2003, 103, 893.

16. Sharma V., Kamal R., Kumar V. Heterocyclic Analogues as Kinase Inhibitors: A Focus Review. Curr. Top Med. Chem. 2017, 17, 2482.

17. Platts M. M. Hydrochlorothiazide, a new oral diuretic. Br. Med. J. 1959, 1, 1565.

18. Martínez A., Esteban A. I., Castro A., Gil C., Conde S., Andrei G., Snoeck R., Balzarini J., De Clercq E. Novel potential agents for human cytomegalovirus infection: synthesis and antiviral activity evaluation of benzothiadiazine dioxide acyclonucleosides. J. Med. Chem. 1999, 42, 1145.

19. Tait A., Luppi A., Hatzelmann A., Fossa P., Mosti L. Synthesis, biological evaluation and molecular modelling studies on benzothiadiazine derivatives as PDE4 selective inhibitors. Bioorg. Med. Chem. 2005, 13, 1393.

20. Ma X., Wei J., Wang C., Gu D., Hu Y., Sheng R. Design, synthesis and biological evaluation of novel benzothiadiazine derivatives as potent PI3Kδ-selective inhibitors for treating B-cell-mediated malignancies. Eur. J. Med. Chem. 2019, 170, 112.

21. Larsen A. P., Francotte P., Frydenvang K., Tapken D., Goffin E., Fraikin P., Caignard D. H., Lestage P., Danober L., Pirotte B., Kastrup J. S. Synthesis and Pharmacology of Mono-, Di-, and Trialkyl-Substituted 7-Chloro-3,4-dihydro-2H-1,2,4-benzothiadiazine 1,1-Dioxides Combined with X-ray Structure Analysis to Understand the Unexpected Structure-Activity Relationship at AMPA Receptors. ACS Chem. Neurosci. 2016, 7, 378.

22. Hirayama F., Koshio H., Katayama N., Ishihara T., Kaizawa H., Taniuchi Y., Sato K., Sakai-Moritani Y., Kaku S., Kurihara H., Kawasaki T., Matsumoto Y., Sakamoto S., Tsukamoto S. Design, synthesis and biological activity of YM-60828 derivatives. Part 2: potent and orally-bioavailable factor Xa inhibitors based on benzothiadiazine-4-one template. Bioorg. Med. Chem. 2003, 11, 367–381.

23. Kamal A., Reddy K.-S., Ahmed S.-K., Khan M.-N., Sinha R.-K., Yadav J.-S., Arora S.-K. Anti-tubercular agents. Part 3. Benzothiadiazine as a novel scaffold for anti-Mycobacterium activity. Bioorg. Med. Chem. 2006, 14, 650–658.

24. Kamal A., Shetti R.-V, Azeeza S., Ahmed S.-K., Swapna P., Reddy A.-M., Khan I.-A., Sharma S., Abdullah S.-T. Anti-tubercular agents. Part 5: synthesis and biological evaluation of benzothiadiazine 1,1-dioxide based congeners. Eur. J. Med. Chem. 2010, 45, 4545–4553.

25. Tait A., Luppi A., Franchini S., Preziosi E., Parenti C., Buccioni M., Marucci G., Leonardi A., Poggesi E., Brasili L. 1,2,4-Benzothiadiazine derivatives as alpha1 and 5-HT1A receptor ligands. Bioorg. Med. Chem. Lett. 2005, 15, 1185–1188.

26. Hu Z., Martí J., Lu H. Structure of benzothiadiazine at zwitterionic phospholipid cell membranes. J. Chem. Phys. 2021, 155, 154303.

27. Jo S., Kim T., Iyer V. G., Im W. CHARMM-GUI: a web-based graphical user interface for CHARMM. J. Comput. Chem. 2008, 29, 1859.

28. Jo S., Lim J. B., Klauda J. B., Im W. CHARMM-GUI Membrane Builder for mixed bilayers and its application to yeast membranes. Biophys. J. 2009, 97, 50.

29. Jorgensen W. L., Chandrasekhar J., Madura J. D., Impey R. W., Klein M. L. Comparison of simple potential functions for simulating liquid water. J. Chem. Phys. 1983, 79, 926.

30. Klauda J. B., Venable R. M., Freites J. A., et al. Update of the CHARMM all-atom additive force field for lipids: validation on six lipid types. J Phys. Chem. B 2010, 114, 7830.

31. Lim J. B., Rogaski B., Klauda J. B. Update of the cholesterol force field parameters in CHARMM. J. Phys. Chem. B 2012, 116, 203.

32. Huang J., MacKerell Jr A.D. CHARMM36 all-atom additive protein force field: Validation based on comparison to NMR data. J. Comput. Chem. 2013, 34, 2135.

33. Linse B., Linse P. Tuning the smooth particle mesh Ewald sum: Application on ionic solutions and dipolar fluids. J. Chem. Phys. 2014, 141, 184114.

34. Heinzinger K. Computer Modelling of Fluids Polymers and Solids (Springer) 1990, pp. 357–394.

35. Brodholt J. P. Molecular dynamics simulations of aqueous NaCl solutions at high pressures and temperatures. Chem. Geol. 1998, 151, 11.

36. Sala J., Guàrdia E., Martí J. Specific ion effects in aqueous eletrolyte solutions confined within graphene sheets at the nanometric scale. Phys. Chem. Chem. Phys. 2012, 14, 10799.

37. Chowdhuri S., Chandra A. Molecular dynamics simulations of aqueous NaCl and KCl solutions: Effects of ion concentration on the single-particle, pair, and collective dynamical properties of ions and water molecules. J. Chem. Phys. 2001, 115, 3732.

38. Joseph S., Mashl R. J., Jakobsson E., Aluru N. R. Electrolytic transport in modified carbon nanotubes. Nano Lett. 2003, 3, 1399.

39. Allen T. W., Andersen O. S., Roux B. Molecular dynamics—potential of mean force calculations as a tool for understanding ion permeation and selectivity in narrow channels. Biophys. Chem. 2006, 124, 251.

40. Yang J., Calero C., Martí J. Diffusion and spectroscopy of water and lipids in fully hydrated dimyristoylphosphatidylcholine bilayer membranes. J. Chem. Phys. 2014, 140, 104901.

41. Lu H., Martí J. Binding and dynamics of melatonin at the interface of phosphatidylcholine-cholesterol membranes PLoS One 2019, 14, e0224624.

42. Drew Bennett W.F., He S., Bilodeau C. L., Jones D., Sun D., Kim H., Allen J. E., Lightstone F. C., Ingólfsson H. I. Predicting small molecule transfer free energies by combining molecular dynamics simulations and deep learning. J. Chem. Inf. Model. 2020, 60, 5375.

43. Lu H., Martí J. Cellular absorption of small molecules: free energy landscapes of melatonin binding at phospholipid membranes. Sci. Rep. 2020, 10, 9235.

44. Abraham M. J., Murtola T., Schulz R., et al. GROMACS: High performance molecular simulations through multi-level parallelism from laptops to supercomputers. SoftwareX 2015, 1, 19.

45. Pronk S., Páll S., Schulz R., et al. GROMACS 4.5: a high-throughput and highly parallel open source molecular simulation toolkit. Bioinformatics 2013, 29, 845.

46. Van Der Spoel D., Lindahl E., Hess B., et al. GROMACS: fast, flexible, and free. J Comput. Chem. 2005, 26, 1701.

47. Lindahl E., Hess B., Van Der Spoel D. GROMACS 3.0: a package for molecular simulation and trajectory analysis. Molecular modeling annual 2001, 7, 306.

48. Berendsen H. J. C., Van der Spoel D., Van Drunen R. GROMACS: A message-passing parallel molecular dynamics implementation. Comput. Phys. Commun. 1995, 91, 43.

49. Chen W., Dusa F.,Witos J.,Ruokonen S.-K., Wiedmer S. K. Determination of the main phase transition temperature of phospholipids by nanoplasmonic sensing. Sci. Rep. 2018, 8, 14815.

50. Evans, D.J., Holian, B.L. The Nose–Hoover thermostat. J. Chem. Phys. 1985, 83, 4069–4074.

51. Parrinello, M., Rahman, A. Crystal structure and pair potentials: A molecular-dynamics study. Phys. Rev. Lett. 1980, 45, 1196.

52. Lu, H., Martí, J. Long-lasting Salt Bridges Provide the Anchoring Mechanism of Oncogenic Kirsten Rat Sarcoma Proteins at Cell Membranes. J. Phys. Chem. Lett. 2020, 11, 9938–9945.

53. Pandey, P.R., Roy, S. Headgroup mediated water insertion into the DPPC bilayer: a molecular dynamics study. J. Phys. Chem. B 2011, 115, 3155–3163.

54. Pan, J., Tristram-Nagle, S., Nagle, J.F. Effect of cholesterol on structural and mechanical properties of membranes depends on lipid chain saturation. Phys. Rev. E 2009, 80, 021931.

55. Martí, J., Lu, H. Microscopic interactions of melatonin, serotonin and tryptophan with zwitterionic phospholipid membranes. Int. J. Molec. Sci. 2021, 22, 2842.

56. Liu, H., Zhang, H., Jin, B. Fluorescence of tryptophan in aqueous solution. Spectrochimica Acta Part A: Molecular and Biomolecular Spectroscopy 2013, 106, 54–59.

57. Trzesniak, D., Kunz, A-P.E., van Gunsteren, W.F. A comparison of methods to compute the potential of mean force. ChemPhysChem 2007, 8, 162–169.

58. Guàrdia, E., Rey, R., Padró, J.A. Statistical errors in constrained molecular dynamics calculations of the mean force potential. Mol. Sim. 1992, 9, 201–211.

59. Kästner, J. Umbrella sampling. Wiley Interdisciplinary Reviews: Comp. Mol. Sci. 2011, 1, 932–942.

60. Geissler, P.L., Dellago, C., Chandler, D., Hutter, J., Parrinello, M. Autoionization in liquid water. Science 2001, 291, 2121–2124.

61. Martí, J., Csajka, F.S., Chandler, D. Stochastic transition pathways in the aqueous sodium chloride dissociation process. Chem.Phys.Lett. 2000, 328, 169–176.

62. Bolhuis, P.G., Chandler, D., Dellago, C., Geissler, P.L. Transition path sampling: Throwing ropes over rough mountain passes, in the dark. Annu.Rev.Phys.Chem. 2002, 53, 291–318.

63. Martí, J., Csajka, F.S. Transition path sampling study of flip-flop transitions in model lipid bilayer membranes. Phys.Rev.E 2004, 69, 061918.

64. Barducci, A., Bussi, G., Parrinello, M. Well-tempered metadynamics: a smoothly converging and tunable free-energy method. Phys.Rev.Lett. 100, 2008, 020603.

65. Chandler, D. Introduction to Modern Statistical Mechanics (Oxford University Press, Oxford, UK) 1987.

66. Feyereisen, M.W., Feller, D., Dixon, D.A. Hydrogen bond energy of the water dimer. J Phys. Chem. 1996, 100, 2993–2997.

67. Vega, C., Abascal, J.L.F., Conde, M.M., Aragones, J.L. What ice can teach us about water interactions: a critical comparison of the performance of different water models. Faraday Discuss. 2009, 141, 251–276.

68. MacCallum, J.L., Bennett, W.F.D., Tieleman, D.P. Distribution of amino acids in a lipid bilayer from computer simulations. Biophys.J. 2008, 94, 3393–3404.

69. de Jesus, A.J., Allen, T.W. The role of tryptophan side chains in membrane protein anchoring and hydrophobic mismatch. BBA. Biomembranes 2013, 1828, 864–876.

70. Peters, G.H., Werge, M., Elf-Lind, M.N., Madsen, J.J., Velardez, G.F., Westh, P. Interaction of neurotransmitters with a phospholipid bilayer: a molecular dynamics study. Chem. and Phys. of lipids 2014, 184, 7–17.

71. Lu, H., Martí, J. Effects of cholesterol on the binding of the precursor neurotransmitter tryptophan to zwitterionic membranes. J. Chem. Phys. 2018, 149, 164906.

72. Lu, H., Martí, J. Cellular absorption of small molecules: free energy landscapes of melatonin binding at phospholipid membranes. Sci.Rep. 2020, 10, 1–12.

73. Martí, J., Padró, J.A., Guàrdia, E. Computer simulation of molecular motions in liquids: Infrared spectra of water and heavy water. Mol.Sim. 1993, 11, 321–336.

74. Padró, J.A., Martí, J. Response to “Comment on ‘An interpretation of the low-frequency spectrum of liquid water’”[J. Chem. Phys. 118, 452 (2003)]. J. Chem. Phys. 2004, 120, 1659–1660.

75. Padró, J.A., Martí Guàrdia, E. Molecular dynamics simulation of liquid water at 523 K. J.Phys.:Condensed Matter 1994, 6, 2283.

76. Martí, J., Gordillo, M.C. Microscopic dynamics of confined supercritical water. Chem.Phys.Lett. 354, 2002, 227–232.

77. Martí, J. Dynamic properties of hydrogen-bonded networks in supercritical water. Phys.Rev.E 2000, 61, 449.

